# DeepMicrobes: taxonomic classification for metagenomics with deep learning

**DOI:** 10.1101/694851

**Authors:** Qiaoxing Liang, Paul W. Bible, Yu Liu, Bin Zou, Lai Wei

## Abstract

Taxonomic classification is a crucial step for metagenomics applications including disease diagnostics, microbiome analyses, and outbreak tracing. Yet it is unknown what deep learning architecture can capture microbial genome-wide features relevant to this task. We report DeepMicrobes (https://github.com/MicrobeLab/DeepMicrobes), a computational framework that can perform large-scale training on > 10,000 RefSeq complete microbial genomes and accurately predict the species-of-origin of whole metagenome shotgun sequencing reads. We show the advantage of DeepMicrobes over state-of-the-art tools in precisely identifying species from microbial community sequencing data. Therefore, DeepMicrobes expands the toolbox of taxonomic classification for metagenomics and enables the development of further deep learning-based bioinformatics algorithms for microbial genomic sequence analysis.

## Introduction

Shotgun metagenomic sequencing provides an unprecedented high-resolution insight into the critical roles of microorganisms in human health and environment^1^. One of the fundamental analysis steps in metagenomics is to assign individual reads to their species-of-origin. Unlike 16S rRNA sequencing data, which ignores more than 99% of the genomic sequences, taxonomic classification of whole genome shotgun sequencing data is more challenging and capacity demanding for machine learning algorithms. The models should learn genome-wide patterns during training, whereas only information from a short genomic fragment is available during application. Current taxonomic classification algorithms mainly utilize handcrafted sequence composition features such as oligonucleotide frequency^2,3^. However, they are either too slow to process large data sets or comparable to, if not worse than, traditional alignment in terms of precision and recall^4^. Additionally, the features used by these models are often too inflexible to meet the requirement of specific applications beyond their original narrow use cases.

Deep learning is a class of machine learning algorithms capable of modeling complex dependencies between input data (e.g., genomic fragments) and target variables (e.g., species-of-origin) in an end-to-end fashion. Thanks to graphical processing units (GPUs), deep learning-based bioinformatics tools can rapidly process large amounts of metagenomics sequencing data. We thus hypothesize that deep learning can automatically discover taxonomic classification-relevant and genome-wide shared features appearing in short metagenomics sequencing reads given a well-designed deep neural network (DNN) architecture.

Deep learning has made tremendous recent advances in genomics^5^. Taking one-hot encoded DNA sequences as input, the DNNs that have been employed to genomic data fall into two major categories, convolutional neural networks (CNNs) and a hybrid of CNNs and recurrent neural networks (RNNs). For example, DeepSEA^6^, PrimateAI^7^ and SpliceAI^8^ used CNNs to predict the impact of genetic variation. Seq2species^9^ also adopted CNNs to predict the species-of-origin of 16S data. DeeperBind^10^ and DanQ^11^ used hybrid architectures to predict transcription factor binding and DNA accessibility. Despite the success of these applications, it remains unknown what DNN architecture and DNA encoding method are suitable for taxonomic classification of metagenomics data.

Here we describe DeepMicrobes, a *k*-mer embedding-based recurrent network with attention mechanism (**Fig. 1a**). We trained the DNN on synthetic reads from RefSeq complete bacterial and archaeal genomes. The first layer of DeepMicrobes is designed to encode *k-*mers to dense vectors through embedding. The vectors are fed into a bidirectional long short-term memory network (BiLSTM) followed by self-attention and a multilayer perceptron (MLP).

**Figure 1.**
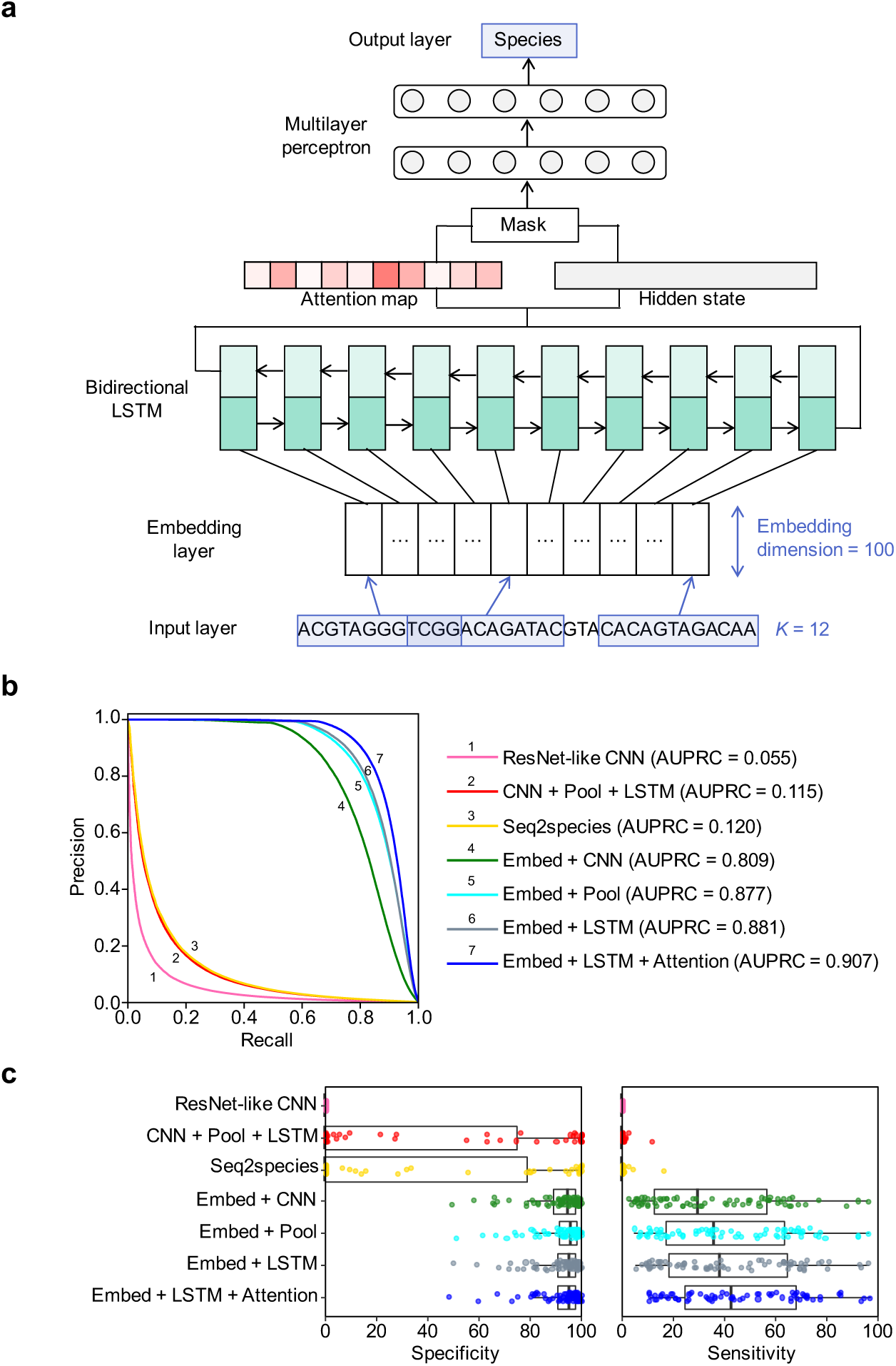
The architecture of DeepMicrobes and the performance of different DNN methods. (**a**) The deep learning architecture of DeepMicrobes. (**b**) The AUPRC of different models on the synthetic test set consists of reads from 1,000 microbial genomes in equal proportion. (**c**) The specificity (left) and sensitivity (right) of different models on the genome sequencing data. ResNet-like CNN, a convolutional model; CNN + Pool + LSTM, a hybrid convolutional and recurrent model; Seq2species, a previously proposed architecture for 16S data; Embed + CNN, an embedding-based convolutional model; Embed + Pool, an embedding baseline; Embed + LSTM, an embedding-based recurrent model; Embedding + LSTM + Attention, an embedding-based recurrent self-attention model, which is selected for DeepMicrobes.

DeepMicrobes surpasses other explored architectures both on synthetic and real sequencing data. Specifically, *k*-mer embedding rather than one-hot encoding boosts model performance. In addition, we show that our deep learning approach produces less false positive identifications than other taxonomic classification tools based on database searching.

## Results

### A deep learning architecture for taxonomic classification

To determine what kind of DNN is suitable for modeling the taxonomic signatures of shotgun metagenomic sequencing reads, we presented a systematic exploration of DNN architectures with different combinations of network architectural building blocks, DNA encoding schemes, and other hyperparameters. We used a curated RefSeq complete bacterial genome subset for model selection to release the computational burden of architecture searching (Methods). The training set consisted of simulated 100 bp reads in equal proportion from each species. To test the performance of these models, we created a synthetic data set consisted of 100,000 read pairs in equal proportion from 1,000 genomes (**Supplementary Table 1**). We used genome sequencing data sets from Sequence Read Archive (SRA) to evaluate their robustness on real data (**Supplementary Table 2-3**). We used confidence > 50% as the threshold for classified reads.

To determine whether the deep learning architectures for other DNA sequence modeling tasks can be transferred to taxonomic classification, we respectively trained the models which are representatives of two major types of previously employed DNNs. We began with ResNet-like convolutional models, which achieved state-of-the-art performance in predicting the impact of mutations^7,8^. The convolutional models took as input one-hot encoded DNA sequences, and fed them into multiple stacking convolutional blocks (Methods). We varied the number of convolutional blocks and found that the model with six blocks achieved the highest area under the precision recall curve (AUPRC = 0.055), followed by eight blocks (AUPRC = 0.052) on the synthetic data set (**Fig. 1b**). Due to low-confidence predictions, the sensitivity and specificity of the model were closed to zero on the real data sets (**Fig. 1c** and **Supplementary Table 2-3**).

We next trained the hybrid architecture of CNN and RNN, which was proved to be effective in predicting transcription factor binding^10,11^. One-hot encoded DNA sequences were fed to a convolutional layer followed by BiLSTM (Methods). Despite its simplicity, the hybrid model (AUPRC = 0.115) surpassed the ResNet-like CNN (**Fig. 1b**). Also, the hybrid model achieved higher than 90% specificity for 16 out of 72 real sequencing data sets (**Fig. 1c** and **Supplementary Table 2**). Nonetheless, the sensitivity remained low due to the low prediction confidence (**Fig. 1c** and **Supplementary Table 3**).

Seq2species was an architecture designed for predicting species-of-origin of 16S data^9^. Taking one-hot encoded DNA vectors as input, seq2species used depthwise separable convolution as its main component. We retrained the model to assess whether this architecture could be transferred to shotgun metagenomic reads classification (Methods). It is worth noting that we used a batch size of 2,048 which performed better than 500 as of training on 16S data, suggesting the importance of larger batch size in metagenomic setting. In general, the performance of seq2species was only slightly better than the hybrid model both on the synthetic data (AUPRC = 0.120) and the real sequencing data (**Fig. 1b, c**). These results demonstrate that applying subtle variants of CNN or combination with RNN provides more performance boost than stacking a deeper CNN for shotgun metagenomic sequences classification.

The deep learning architectures above share the idea that a convolutional layer should be adopted as the first layers to locate pattern features from one-hot encoded DNA sequences. Indeed, CNNs excel in the recognition of motifs, which is helpful for predicting splicing site and *cis*-acting elements like promoter and enhancer. Notably, CNNs might not take into account the spatial ordering of local motifs, given the evidence from image classification^12^. This can have little impact on tasks where only the occurrence of a few nucleotides in a DNA sequence are the key to classification (e.g. transcription factor binding site detection). However, it is more complex to model taxonomic signatures, such as single-nucleotide variants (SNVs), insertion–deletions (indels), and unique genes, especially for short microbial sequencing reads. Moreover, one-hot encoding has its own limitations. Apart from being sparse in information, one-hot approach encodes double strands of a DNA sequence into two unrelated matrices.

To overcome these limitations, we made an analogy between *k*-mers and words and used *k*-mer embedding to represent DNA sequences, which is common practice in natural language processing (NLP). Reverse complement *k*-mers are treated as the same word (Methods). To determine the contribution of this encoding scheme to model performance, we trained an embedding-based baseline model whose only trainable parameters were the weights in the embedding layer (Methods). The preliminary experiments showed that the models performed better using longer *k*-mer, thus we chose the longest *k*-mer (*k* = 12) whose vocabulary was able to fit in the memory of our GPUs. Interestingly, the baseline model (AUPRC = 0.877) outperformed the models taking one-hot encoded DNA as input (**Fig. 1b**). On the real sequencing data sets, the model assigned reads to the target species in > 90% specificity for 56 data sets, and > 95% specificity for 39 data sets (**Fig. 1c** and **Supplementary Table 2**). Meanwhile, all the target species was successfully identified (**Fig. 1c** and **Supplementary Table 3**). This implies that the *k*-mer embedding layer is capable of embedding taxonomic attributes in each *k*-mer vector.

We next asked what types of neural networks were appropriate to learn useful information from *k*-mer embedding. To investigate this, we made two extensions on the baseline model by respectively adding a convolutional and BiLSTM layer after the *k*-mer embedding layer (Methods). Surprisingly, the embedding-based convolutional model was worse than the baseline model on the synthetic data (AUPRC = 0.809) and real genomic sequencing data in specificity (*P* < 6.7 × 10^-5^) and sensitivity (*P* < 1.8 × 10^-22^), though it contained more parameters in the convolutional layer (**Fig. 1b, c**). In contrast, the embedding-based recurrent model (AUPRC = 0.881) further increased the performance of the baseline on the real data in specificity (*P* < 1.1 × 10^-2^) and sensitivity (*P* < 1.9 × 10^-4^; **Fig. 1b, c**).

Self-attention is an attention mechanism capable of extracting relevant aspects from sentences with no need for additional information^13^. Inspired by its successful applications in a variety of NLP tasks, we applied self-attention mechanism on top of the BiLSTM of the embedding-based recurrent model to evaluate if this could further improve the model performance (Methods). Instead of directly taking the hidden state of the BiLSTM as features for the MLP, self-attention enabled the model to focus on specific regions of input DNA sequences, and generated sequence-level representation. Indeed, the model reached an AUPRC of 0.907 (**Fig. 1b**). When evaluated on the real sequencing data sets, the model also surpassed the embedding-based recurrent model without self-attention mechanism in specificity (*P* < 1.2 × 10^-2^) and sensitivity (*P* < 2.6 × 10^-14^; **Fig. 1c**). This suggests that paying more attention to some specific parts of reads might help DNNs better model the unique features among short genomic sequences from different microorganisms. This deep learning architecture is selected and termed DeepMicrobes in the following studies (**Fig. 1a**).

To confirm the impact of embedding *k*-mer length, we trained a series of variant models of DeepMicrobes using *k* < 12. We observed that on the synthetic data the AUPRC increased from 0.255 (*k* = 8) to 0.589 (*k* =11), and the trend was consistent on the real data (**Supplementary Fig. 1** and **Supplementary Table 4-5**). These results support the potential of using even longer *k*-mer to improve the performance. Other architectures, such as hierarchical attention networks^14^ and the Transformer^15^ that entirely based on attention mechanisms, have potential in taking more advantage of the information in *k*-mer embedding. But they were too big to be trained on shotgun metagenomic reads classification task, by hindering the use of large batch sizes. Taken together, DeepMicrobes is the best feasible deep learning architecture in our problem setting.

### DeepMicrobes generalizes to different taxonomic ranks and read lengths

To test the general applicability of DeepMicrobes on different taxonomic ranks, we used the same architecture as species-level model to build the classifiers at the level of genus, family, order, class and phylum, respectively (Methods). We evaluated the six models on the synthetic data sets whose read lengths was different from the 100 bp training sets. As expected, when tested on the 100 bp data, the performance consistently increased from species to phylum, reaching an AUPRC of 0.951 at the genus level, and a nearly perfect AUPRC of 0.998 at the phylum level (**Fig. 2a**). This probably resulted from reduced burden to the models in distinguishing similar taxa. The monotonically increasing pattern with taxonomic ranks retained for 150 bp and 200 bp test sets, while the 250 bp and 300 bp test sets showed small fluctuation of AUPRC between 0.989 and 0.995 (**Fig. 2a**). The species-level model performed better on longer sequences, with an AUPRC of 0.974 on the 150 bp data set (**Fig. 2a**). Interestingly, the models at the level higher than order tended to perform better on the read lengths similar to training sets. Nonetheless, the AUPRCs of these high-rank models were all above 0.99. These results indicate the overall robustness of DeepMicrobes on multiple taxonomic ranks and varying length of reads that were not seen during training.

**Figure 2.**
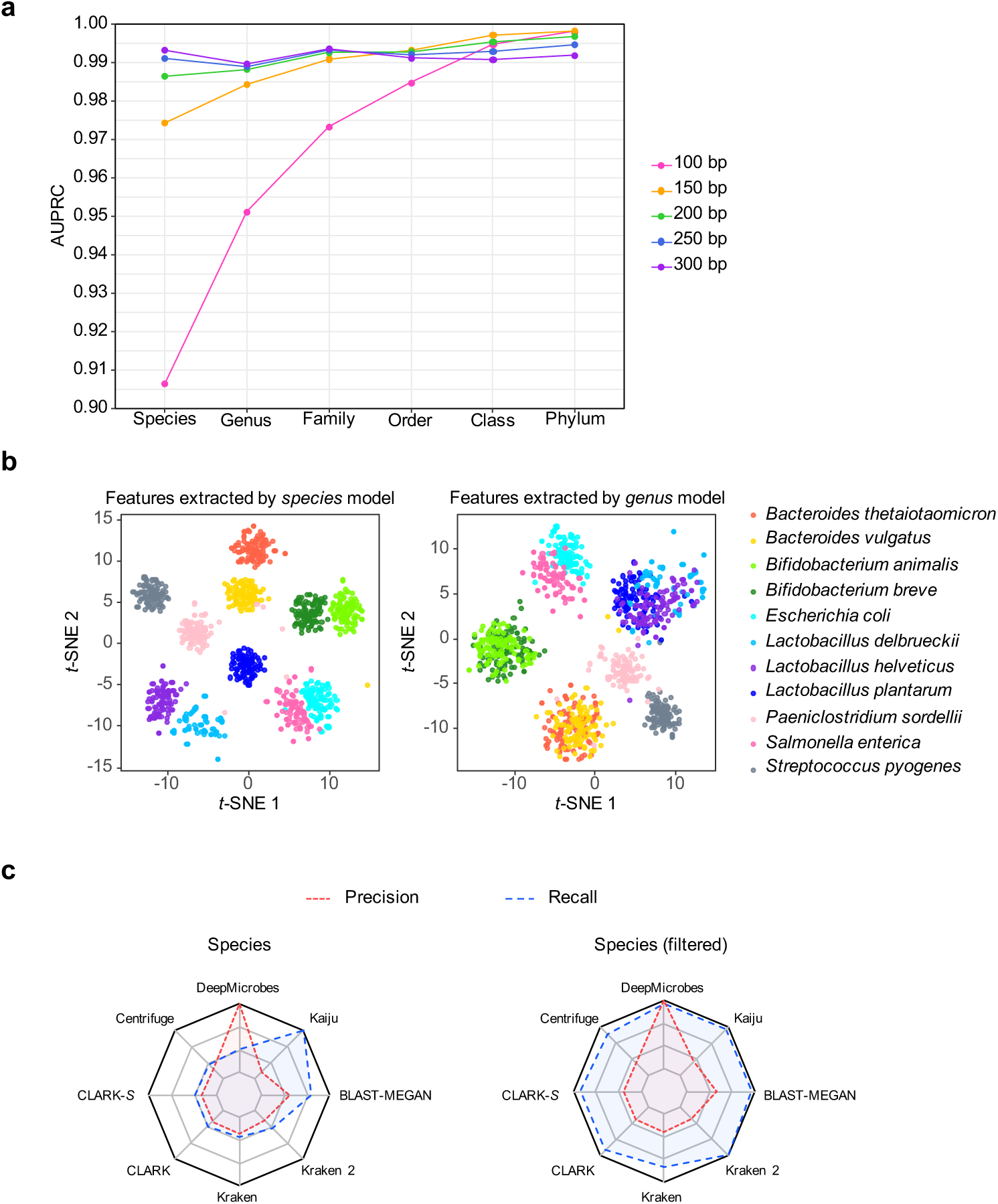
Generalization of DeepMicrobes to different taxonomic ranks, and comparison of DeepMicrobes with state-of-the-art tools. (**a**) The test performance of DeepMicrobes taxonomic-rank variants on reads of different lengths. We used the synthetic test sets containing reads from 1,000 genomes in equal proportion. Each model variant was trained on 100 bp reads, and tested on 100 bp (magenta), 150 bp (orange), 200 bp (green), 250 bp (blue), and 300 bp (purple) reads. (**b**) T-SNE visualization of the mock community reads using high-level feature maps generated by DeepMicrobes trained at the species (left) and genus (level) level. (**c**) Relative precision and recall of the medium-complexity CAMI data set at the species level, computed on the basis of default (left) and shared-species filtered (right) results. Metrics were normalized by the maximal value.

Unlike traditional species classification approaches based on alignment, *k*- mer frequency or machine learning systems with hand engineered features, our deep neural network extracts novel, useful, and reusable features from the underlying data sets. We hypothesized that DeepMicrobes makes robust predictions by extracting high-level features that are shared among hundreds-of-nucleotide fragments across the genomes of a taxon from primary metagenomic sequences. To test this hypothesis, we used a published mock community sample consisted of 11 species members from 7 genera^16^, and obtained the feature maps generated from the last hidden layer of the species-level model and genus-level model, respectively (Methods).

We then applied t-Distributed stochastic neighbor embedding (T-SNE) dimension reduction^17^ to visualize a randomly drawn subset of the metagenome sample using these features (Methods). The sequencing reads originated from the same species clustered into unique groups (**Fig. 2b**). Furthermore, the distance between clusters could partly reflect the evolutionary relationships. Species of the same genus tended to be closer (**Fig. 2b**). *Escherichia coli* and *Salmonella enterica*, reported to share a supraspecies pangenome^18^, also showed this pattern (**Fig. 2b**). When using features extracted by the genus-level model, species of the same genus mixed together to form one big cluster (**Fig. 2b**). This pattern indicates that the characteristics of the learned features depend on the training target allowing for a flexible and tunable approach to extracting meaningful sequence features. Thus, DeepMicrobes could potentially extract more specific features that enable discrimination among even more similar taxa such as strain provided suitable training data is available. Notably, one of the species (and also genus) in the community, *Paeniclostridium sordellii*, was excluded from the training sets due to incomplete genomes. Nonetheless, the taxon formed a distinguishable cluster both at the species and genus level (**Fig. 2b**), demonstrating the versatility of the high-level features in grouping microbial sequences from the same taxon as well as identifying novel organisms that were not part of the training data.

### DeepMicrobes improves species identification by database searching

Accurate species identification from metagenomics samples is a critical aspect of taxonomic classification. However, most database search tools for metagenomics only retain high precision and recall until the family level^19^. To test the advantage of our deep learning-based approach in species and genus identification, we analysed the Critical Assessment of Metagenome Interpretation (CAMI) data sets^19^ using DeepMicrobes and seven other taxonomic classification tools, including Kraken^20^, Kraken 2, Centrifuge^21^, CLARK^22^, CLARK-*S*^23^, Kaiju^24^ and BLAST-MEGAN^25^. To this end, we trained DeepMicrobes to assign species label to reads using 10,857 RefSeq complete bacterial and archaeal genomes covering 3,640 species (Methods). Apart from the classification results generated using their pre-built reference databases if available, we also filtered the results to only consider the species shared by all reference databases or training set. This was to eliminate the effect of database setting on performance metrics, and focus on the algorithms.

We observed that DeepMicrobes substantially outperformed other tools in terms of precision at the species and genus level (**Fig. 2c** and **Supplementary Fig. 2**). For example, Kraken identified 1,754 more false positive species than DeepMicrobes from the medium-complexity sample, based on the filtered results. Specifically, DeepMicrobes identified less false positive species than BLAST-MEGAN regardless of the database setting. Meanwhile, DeepMicrobes classified reads faster than the other tools, except for Kraken and Kraken 2 (**Supplementary Fig. 2**). In detail, when processing 100 bp reads, DeepMicrobes was 1.3 times faster than Centrifuge, and 519.9 times faster than BLAST-MEGAN, which was the second most precise tool. Notably, we used eight CPUs to run the other tools and one GPU to run DeepMicrobes. Moreover, since the number of reads that can be processed in parallel totally relies on available memory, the classification speed of DeepMicrobes has large room to improve given a more powerful GPU than the one used in this study. These results suggest that our deep learning approach has advantages over database searching, especially when false positives would strongly increase the cost in downstream efforts.

## Discussion

In this study, we introduce DeepMicrobes, a deep learning architecture able to accurately predict the species-of-origin from primary shotgun metagenomic sequencing reads. Although trained on simulated reads, it performed well on real data with sequences different from the genomes used for training. Including real sequencing reads might further improve the performance.

Effective taxonomic classification requires a distinct DNA encoding scheme and deep neural network architecture for precise genomic modeling tasks. We show that replacing one-hot encoding with *k*-mer embedding significantly boosts model performance. One likely reason for the improvement may be that taxonomic information is encoded by the *k*-mer representations in the embedding space. With this representation, difference between similar sequences originating from closely related species could be amplified. Finally, a pair of reverse complement DNA sequences consist of the same words, thus knowledge could be easily transferred between them. Interestingly, *k*-mer embedding has recently been showed to surpass one-hot encoding in predicting transcription factor binding^26^. This suggests the general applicability of *k*-mer embedding in other biological fields.

Our finding that RNNs surpass CNNs in taxonomic classification highlights the importance of order and context of oligonucleotides in taxonomic classification. CNNs and *k*-mer exact alignments only take into account the presence of specific oligonucleotides, while early machine learning-based taxonomic classifiers employed their frequency as features. In contrast, BiLSTM understands a *k*-mer better with the help of knowledge learned from the previous and next *k*-mer. Hence, ordering and contextual information are retained and passed to the next layer.

To our knowledge, DeepMicrobes is the first deep learning architecture that incorporates attention mechanisms in DNA sequences analysis. Apart from a performance boost, attention scores provide an easy way to visualize what parts of the DNA sequences contribute most to prediction. This characteristic makes algorithm more interpretable than perturbation-based and backpropagation-based approaches opening the possibility of exploring the biological meaning of the extracted features in contrast to black-box prediction algorithms.

DeepMicrobes provides a novel tool and information source for taxonomy identification and expands the repertoire of metagenome analysis methods. Unlike algorithms based on read mapping, discriminative *k*-mer, or sequence composition, DeepMicrobes extracts task-relevant features from DNA sequences using a deep neural network learning architecture. Notably, the *k*- mer length we recommend is far from being discriminative among species as is the case of Kraken, CLARK, and Centrifuge. Current binning methods using sequence compositions as features typically perform well on long contigs. However, we show that the sequences as short as 100 bp formed separable clusters using high-level features extracted by DNNs. The feature type generated by supervised learning depends on training targets, which is more focused and task-relevant than auto-encoder methods^27^. Future researches might investigate how to utilize these features, and also incorporate them with co-occurrence or coverage information to build a powerful metagenome binning tool.

We demonstrate that DeepMicrobes is capable of discovering microbial genome-wide features appearing in short genomic fragments. Apart from general microbiome analysis, taxonomic classification is also useful in other scenarios such as outbreak tracing, pathogen identification, and virulence prediction. Given the flexibility and expressiveness of deep learning modeling techniques, DeepMicrobes might be easily transferred to these tasks by shifting training sets. For example, predicting the source of food-borne disease would require the deep learning model to be trained on whole-genome sequencing data of *Salmonella enterica* collected from different hosts^28^. We believe that DeepMicrobes will be of benefit for development of deep learning-based bioinformatics tools that are able to extract new insights from the exponentially increasing amount of microbial genomic sequencing data.

## Methods

### Data sets for model training

Source genomes for training were collected from National Center for Biotechnology Information (NCBI) reference sequences (RefSeq) bacterial and archaeal genome database (downloaded on 2018-11-30). Training sets were constructed by simulating sequencing reads from complete genomes using wgsim in the SAMtools software package^29^. We simulated 100 bp error-free reads in equal proportion for each species. The number of reads to simulate depended on how many training steps were required for models to converge. Apart from sampling from both strands of genomes, we also included the reverse complement reads in the training set. Each read was given a numerical label based on NCBI taxonomy IDs at the species level (more details provided with the source code at https://github.com/MicrobeLab/DeepMicrobes). The reads for training at other taxonomic ranks (phylum, class, order, family, and genus) were labeled at that specific rank.

The species included in the training set for model selection was required to contain at least one genome of a strain at the reference or representative assembly level of quality in the RefSeq database. We drew one genome as representative for each species. This training set was also used to train the variants of DeepMicrobes at different taxonomic level (https://github.com/MicrobeLab/DeepMicrobes). The full training set used to train the model for comparison with other taxonomic classifiers contained filtered RefSeq bacterial and archaeal species. We first screened the similar pairs of species using the tetranucleotide signature correlation index implemented by pyani^30^ (http://widdowquinn.github.io/pyani). We next computed the average nucleotide identity (ANI) between these species pairs whose tetranucleotide signature correlation index > 0.99 ^31^ using a window size of 100 bp. If > 80% coverage of the genomes of a species showed an ANI > 95%, the species was excluded from training. This resulted in 10,857 genomes of 3,640 species. Full list of the genomes and species is available at https://github.com/MicrobeLab/DeepMicrobes.

### Data sets for model selection

We created an evaluation set consisting of 10,000 100 bp reads for each species, which was simulated using wgsim with a random seed different from the one used to generate the training sets. This evaluation set was used to search for optimal hyperparameters and decide when to stop training. This data set was not seen during training to protect against overfitting the model. We used Mason read simulator^32^ to create the synthetic test set from 1,000 bacterial genomes (**Supplementary Table 1**). Before genome fragmentation, a SNP rate of 0.1% and an indel rate of 0.1% were injected in genomes. In addition, an indel rate of 0.1% and a mutation rate of 0.4% were injected in the reads. We simulated equal proportion of 100 bp read pairs for each genome, and benchmarked the models with different architectures on this data set with 100,000 reads. We also created the data sets in 150 bp, 200 bp, 250 bp, and 300 bp, which are common lengths for next-generation sequencing reads. The data sets used to evaluate model performance at different taxonomic ranks were the same, except for the true labels were given at the target rank. We downloaded 72 bacterial genome sequencing samples from the Sequence Read Archive (SRA) at NCBI (**Supplementary Table 2**). We filtered reads shorter than 100 bp after quality control, and truncated longer reads to 100 bp.

### Representation of DNA sequences

We adopted two strategies to encode DNA sequences into numeric matrices, namely one-hot encoding and *k*-mer embedding. For one-hot encoding we converted DNA into 4 × *L* matrix, where A = [1, 0, 0, 0], C = [0, 1, 0, 0], G = [0, 0, 1, 0] and T = [0, 0, 0, 1]. For *k*-mer embedding, we split a DNA sequence of length *L* into a list of substrings of length *K* with a stride of *S*. We used a stride of one for the final model, ending up with *L* – *K* + 1 substrings. The length of *K* was chosen to reach balance between the model’s fitting capacity and computational resources since the vocabulary size grows exponentially in *K* by 4*^K^* (**Supplementary Table 6**). We used 12-mers unless otherwise stated. The *k*-mer vocabulary was constructed using Jellyfish. We only retained canonical *k*-mers as representatives (-C parameter of Jellyfish), which downsized the vocabulary (**Supplementary Table 6**). We included a word symbol <UNK> in the vocabulary to represent *k*-mers with Ns. Each *k*-mer was further encoded as a zero-based integer according to its lexical order in the vocabulary. These integers then served as indexes for the word embedding layer.

### Model architectures

#### Convolutional model

The ResNet^33^-like CNN took as input the one-hot encoded DNA sequences. The architecture started with one layer of convolutions, followed by a stack of shortcut connected ResNet-like temporal convolutional blocks. One convolutional block consisted of two convolutional layers, each followed by a layer of batch normalization and activation. A pooling layer was inserted every two convolutional blocks. This resulted in a DNN with 1 + 2*N* convolutional layers, where *N* is the number of convolutional blocks. Unless otherwise stated, we used MLP as a classifier for species label prediction, which was also the case of the other models.

#### Hybrid convolutional and recurrent model

DNA sequences were input as one-hot matrices. The models began with one convolutional layer and one pooling layer, followed by BiLSTM.

#### Seq2species

We used the hyperparameters of Seq2species optimized for 100 bp reads^9^, except that the number of nodes in the output layer was changed to the number of species in our setting. To train the model, we used the code available at https://github.com/tensorflow/models/tree/master/research/seq2species. To benchmark running time and other performance metrics, we adapted the code to our input and output pipelines without changing the code related to the model architecture (https://github.com/MicrobeLab/DeepMicrobes).

#### Embedding baseline

The *k*-mer embedding layer learned a mapping from each *k*-mer index to an embedding vector. We randomly initialized the parameters in the embedding layer. Before the fully connected layers, we performed max pooling and average pooling over the dimension of token length of the embedding matrix and concatenated together the two feature vectors.

#### Embedding-based convolutional model

We extended the embedding baseline model by adding a convolutional layer after the embedding layer. The 1D convolution kernel was convolved with the input embedding matrix over the dimension of token length. In addition to convolutional layer with one fixed filter width, the feature maps generated by convolutional layers with multiple filter widths could also be concatenated. We optionally applied an over-time pooling over the features before feeding them to the MLP.

#### Embedding-based recurrent model

We applied a BiLSTM over the embedding vector of each *k*-mer. Similar to the convolution extension, we also tried different types of pooling operation over the hidden states generated by the BiLSTM. Alternatively, the hidden states were directly fed to the MLP.

#### Embedding-based recurrent self-attention model

The summation vectors generated by the self-attention operation were used to weight LSTM hidden states. The attention vector was softmax normalized so as to ensure all the weights summed up to one. Multiple rows of attention were used to focus on multiple aspects of the DNA sequences that reflected taxonomic signatures. For downstream classification task, the self-attention weighted hidden states were fed to the MLP.

### Model training and evaluation metrics

The DNNs were implemented using TensorFlow framework. We used NVIDIA Tesla P40 24GB GPU to accelerate computation. The training set was only seen by the models for one time (i.e., epoch = 1). We trained the models till they converged on the evaluation set. Reads in fasta or fastq format were converted to the TensorFlow format TFRecord before loading into the models.

For each architecture of DNN, we performed random search to pick the optimal combination of hyperparameters. In detail, we randomly sampled 30 candidate hyperparameters setting from the search space (**Supplementary Table 7**) and picked the models which performed best on the evaluation set. We used micro-averaging AUPRC to evaluate model performance on the synthetic test sets. Sensitivity and specificity were used to measure the performance of models on the genome sequencing data sets. Here sensitivity is defined as the proportion of correctly classified reads out of the total number of reads in the sample, and specificity is defined as the proportion of correctly classified reads among all reads classified. The statistical difference was measured by paired t-test.

### Comparison of DeepMicrobes with other taxonomic classifiers

We compared the performance of DeepMicrobes with Kraken, Kraken 2, Centrifuge, CLARK, CLARK-*S*, Kaiju and BLAST-MEGAN, using the CAMI data sets. These tools were run with default options. The tools were run in paired-end mode, except for BLAST-MEGAN. For paired-end data we averaged the softmax probability distributions generated by DeepMicrobes for two ends of reads. We ran Kraken (v1.0) using the pre-built MiniKraken 8GB database included complete bacterial, archaeal, and viral genomes in RefSeq (as of Oct. 18, 2017). We ran Kraken 2 (v2.0.6) using pre-built MiniKraken2 v1 8GB database including RefSeq bacteria, archaea, and viral libraries (available on Apr. 23, 2019). Centrifuge (v1.0.3) was run using pre-built reference database, which was compressed prokaryotic database containing bacteria and archaea (updated on Apr. 15, 2018). The bacteria (and archaea) database for CLARK and CLARK-*S* (v1.2.5) was downloaded via the set_targets.sh script (on Aug. 25, 2018). Kaiju (v1.5.0) was run using pre-built microbial subset of the NCBI nr database (as of May. 16, 2017). To run MEGAN, we first queried unpaired reads using BLAST executable (v2.6.0+) against nt index downloaded from NCBI (on Aug. 25, 2018). We used the Megablast mode and an e-value of 1e-20. Next, we ran MEGAN (V5.3.11) to summarize the lowest common ancestor (LCA) taxon for each read.

Speed was evaluated using 8 threads on the same computer. DeepMicrobes was run with 8 threads on CPU for input pipeline, and 1 GPU for prediction using a batch size of 20,000. The data used for speed evaluation has 1,000,000 reads in 100 bp. We computed the precision and recall for species and genus identification for each tool, demanding at least one supporting reads for the presence of a taxon. Precision refers to the fraction of taxon identified by an analysis tool that is actually present. Recall refers to the fraction of expected taxon that is identified by a tool. The reads whose prediction confidence > 50% were treated as being classified at the species level. Reads with confidence > 45% were treated as classified at the genus of that species.

### Reads clustering using high-level features extracted by DNN

We downloaded the mock community sequencing sample from SRA using accession SRR2081071. The identity of each read was confirmed by running BLAST against nt database. For each species included in training, we randomly sampled 100 reads that were correctly classified by DeepMicrobes. For the species not included we randomly sampled 100 reads from those confirmed via BLAST. We used T-SNE to visualize the feature map generated by the last hidden layer of MLP. Before running T-SNE, we used principal component analysis (PCA) to reduce the features into 150 dimensions explaining > 90% of the variation.

## Code availability

The DeepMicrobes program, trained model parameters, hyperparameters and the implementation of the other DNN architectures are provided at GitHub (https://github.com/MicrobeLab/DeepMicrobes).

## Acknowledgements

We thank all members of the Wei Laboratory for their support and discussion. This work was supported by the National Basic Research Program of China 2015CB964601 and the National Natural Science Foundation of China 81570828.

## Author contributions

L.W. conceived the study; Q.L. and L.W. designed the research; Q.L., P.W.B., Y.L., and B.Z. performed the research; Q.L. and Y.L. analyzed data; P.W.B. and B.Z. contributed analytic tools; Q.L. and P.W.B. drafted the manuscript. All authors read and approved the final version of the manuscript.

---

**Supplementary Figure 1.**
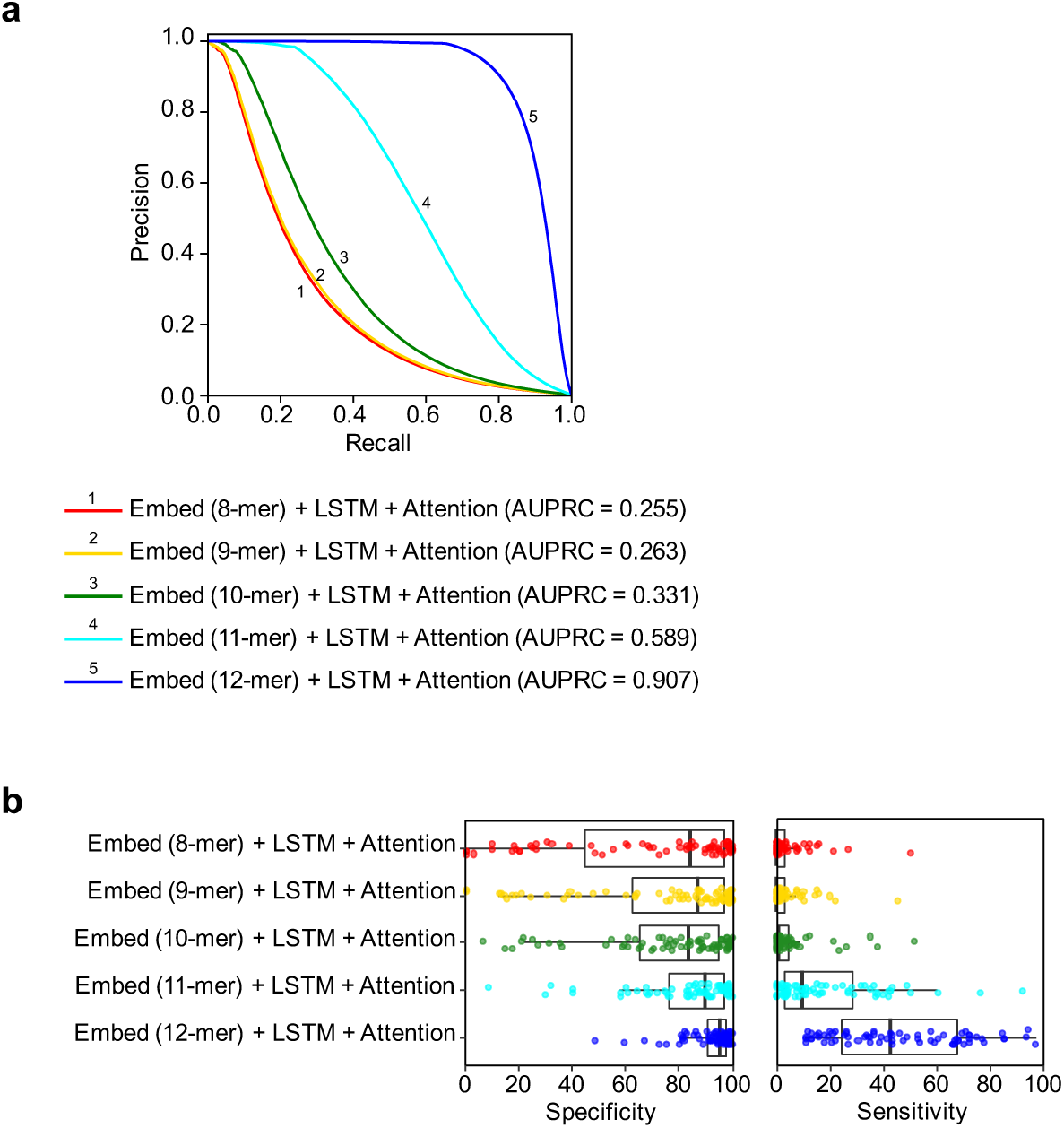
The effect of *k*-mer length on model performance. (**a**) The AUPRC of DeepMicrobes variants using different *k*-mer lengths. The AUPRC was computed on the synthetic test set consisting of reads from 1,000 microbial genomes in equal proportion. The DNA sequences were split into a list of 8-mer, 9-mer, 10-mer, 11-mer and 12- mer, respectively. All model hyperparameters were the same except for the vocabulary size. (**b**) The specificity (left) and sensitivity (right) of DeepMicrobes *k*-mer variants on the genome sequencing data.

**Supplementary Figure 2.**
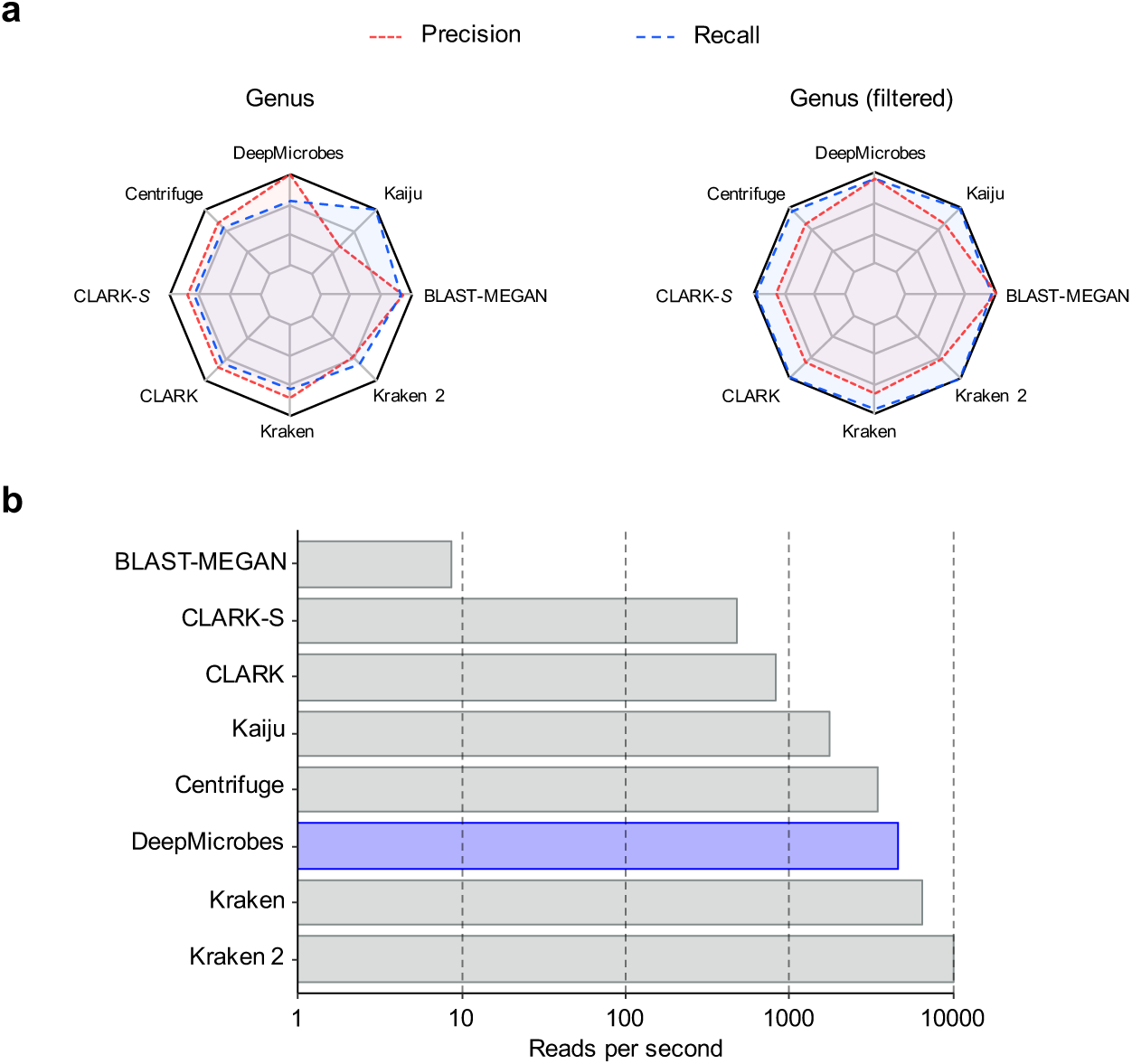
The genus-level comparison of DeepMicrobes with state-of-the-art tools and speed evaluation. (**a**) Relative precision and recall of the medium-complexity CAMI data set at the genus level, computed on the basis of default (left) and shared-genus filtered (right) results. Metrics were normalized by the maximal value. (**b**) Speed comparison of classification programs for 1,000,000 single-end 100 bp reads. DeepMicrobes was run using a batch size of 20,000. The time of DeepMicrobes included converting fasta sequences to TFRecord and making predictions.

Supplementary Table 1. Assembly summary of the 1,000 genomes used to create the synthetic test set

Supplementary Table 2. Specificity of different deep learning architectures on the genome sequencing data set

Supplementary Table 3. Sensitivity of different deep learning architectures on the genome sequencing data set

Supplementary Table 4. Specificity of DeepMicrobes *k*-mer variants on the genome sequencing data set

Supplementary Table 5. Sensitivity of DeepMicrobes *k*-mer variants on the genome sequencing data set

**Supplementary Table 6.** The effect of *k*-mer length on vocabulary size

| Length of *k*-mers | # of all possible *k*-mers ($4^k$) | # of merged *k*-mers (vocabulary size) |
| --- | --- | --- |
| 8 | 65,536 | 32,896 |
| 9 | 262,144 | 131,072 |
| 10 | 1,048,576 | 524,800 |
| 11 | 4,194,304 | 2,097,152 |
| 12 | 16,777,216 | 8,390,656 |

**Supplementary Table 7.** The search space of hyperparameters

| Hyperparameters | Search space |
| --- | --- |
| Number of CNN filters | 64, 128, 256, 320, 512, 1024 |
| Size of CNN filters | 3, 4, 5, 6, 13, 26, 30, concatenate |
| Number of residual block | 1, 2, 3, 4 |
| LSTM dimension (in each direction) | 256, 300, 320, 400, 512, 600, 640, 1024 |
| Number of LSTM layers | 1, 2 |
| Number of FC layers | 1, 2, 3 |
| Number of FC units | 150, 350, 512, 1024, 2048, 3000, 4000, 4096 |
| Type of pooling | Max, average, concatenate, none |
| Window size of pooling | 2, 13, 15, length of input sequence (for embedding models) |
| Pooling stride | 13, 15 |
| Number of attention rows | 10, 20, 30, 40, 50 |
| Penalization coefficient | 0, 1e-5, 1e-4, 1e-3, 0.01, 0.1, 0.2, 0.5, 0.8, 1 |
| Batch size | 128, 256, 500, 512, 1024, 2048, 3000, 4000, 4096 |
| Learning rate | 0.05, 0.01, 5e-3, 1e-3, 5e-4, 1e-4 |
| Decay rate (for learning rate) | 1e-3, 5e-3, 0.01, 0.05, 0.1, 0.5 |
| Dropout (keep probability) | 0.5, 0.6, 0.7, 0.8, 0.9, 0.95, 0.99, 1.0 |
| L2 regularization | 1e-5, 1e-4, 1e-3, 0.01, 0.1, none |
| Activation function | ReLu, tanh, Leaky Relu |
| Optimizer | Adam, Adagrad |
| Encoding method | One-hot, $k$-mer embedding |
| $K$-mer length | 7, 8, 9, 10, 11, 12 |
| $K$-mer redundancy | $4^k$, merged |
| Embedding dimension | 50, 100, 200, 300 |
| Embedding stride | 1, 2, 3 |
| Embedding weights initialization | Random, pre-trained GloVe vectors |

